# Sex-specific organization and synaptic signaling in prefrontal-hypothalamic circuitry

**DOI:** 10.64898/2026.04.29.721673

**Authors:** Courtney A. Bouchet, Emma C. Pinsinski, Julia C. Cook, Chris E. Vaaga, Brent Myers

## Abstract

Top down signaling from the cortex to the hypothalamus is critical to link cognitive and emotional processing to homeostasis and motivation. This study investigates signaling from the medial prefrontal cortex (mPFC) to the posterior hypothalamus (PH), a region that modulates endocrine and autonomic stress responses and motivated behaviors. The function and anatomy of this circuit was examined with patch clamp electrophysiology and mapping studies in male and female rats. Spontaneous firing properties of PH neurons were determined in a cell-type specific manner by combining a transgenic glutamic acid decarboxylase-Cre rat with Cre-dependent colorswitch virus to determine postsynaptic cell-type identity. Overall, PH neurons were more excitable in females compared to males and, in both sexes, data indicated tonic inhibition within the PH, with significantly greater inhibition in males. Using Channelrhodopsin-assisted circuit mapping to query the mPFC-PH circuit, we found that a majority of PH neurons received input from the mPFC and mPFC synapses targeted glutamatergic cells over GABAergic PH cells. Retrograde tracing revealed more PH-projecting neurons in females, specifically within the tenia tecta and infralimbic regions of the mPFC, with significantly more stress-activated PH-projecting cells in the female prelimbic cortex. Anterograde tracing revealed, surprisingly, no sex differences in mPFC presynaptic terminal density in the PH, despite more PH-projecting cell bodies in the female mPFC. These data help to elucidate the sexual divergence in cortical-hypothalamic signaling and how cognitive and emotional information from the prefrontal cortex may differentially regulate homeostasis and motivation between sexes.

**Significance Statement:** Neural signaling between the prefrontal cortex and the hypothalamus is important for maintaining homeostasis, particularly during contextual challenges such as stressors. Here we find multiple aspects of sex-specific organization and neurophysiology in this circuitry. Excitatory inputs from the medial prefrontal cortex target both excitatory and inhibitory neurons within the posterior hypothalamic nucleus in both sexes. However, there are sex differences in the number of stress-activated neurons in the prefrontal cortex that innervate the posterior hypothalamus, as well as differences in hypothalamic inhibitory signaling and estrous cycle-dependent effects on neuronal excitability. Altogether, these data suggest that organizational, synaptic, and hormonal factors may contribute to sex-specific behavioral and physiological integration.

## Introduction

Neural signaling from the medial prefrontal cortex (mPFC) to the hypothalamus is critical to connect higher-order context processing and emotion with homeostatic physiology, but the underlying signaling is not well understood. Additionally, our current understanding of this circuitry is largely derived from male physiology, despite evidence that females are differentially susceptible to mood disorders and physiological co-morbidities (Möller-Leimkühler, 2010). Therefore, understanding cortical-hypothalamic signaling in both sexes can provide neurobiological links between emotional states and physiological outcomes.

Cortical-hypothalamic signaling is important for a variety of behavioral and physiological processes. Signaling from the mPFC to the preoptic and lateral hypothalamus (LH) initiates sleep-preparatory behaviors (Tossell et al., 2023). mPFC projections to the LH also promote social dominance (Padilla-Coreano et al., 2022) and regulate cue-evoked high-fat diet overconsumption (Xiang et al., 2026). Additionally, prefrontal-hypothalamic signaling translates social cues into physiological stress responses. Specifically, signaling from the dorsal tenia tecta / dorsal peduncular cortex to the dorsomedial hypothalamus (DMH) is necessary and sufficient for sympathetic stress responses to social defeat in male rats (Kataoka et al., 2020). Cortical outputs target multiple hypothalamic nuclei that impact stress responses and likely operate in conjunction to orchestrate stress responding (Schaeuble and Myers, 2022). *In vivo* activation of the infralimbic (IL) region of the mPFC has sexually divergent effects on behavioral, endocrine, and cardiovascular responses to stress (Wallace et al., 2021), potentially through robust projections to the posterior hypothalamus (PH) (Wood et al., 2019). The PH is a hypothalamic nucleus that stimulates sympathetic processes including increased body temperature, heart rate, and blood pressure (DiMicco et al., 1986; Lisa et al., 1989), as well as social avoidance, aggression, and anxiety-related behaviors (Myers et al., 2016). In addition to receiving stress-activated inputs from the mPFC (Myers et al., 2016), stress-activated PH neurons send glutamatergic projections to the paraventricular hypothalamic nucleus (PVN) and pre-sympathetic brainstem (Ulrich-Lai et al., 2011; Myers et al., 2016; Nyhuis et al., 2016). Therefore, the PH is anatomically positioned to be a conduit between cortical signaling and physiological stress responses (Myers et al., 2016; Nyhuis et al., 2016; Schaeuble and Myers, 2022). The PH is primarily composed of glutamatergic and GABAergic neurons with a small subset of dopaminergic neurons (Abrahamson and Moore, 2001). In males, excitatory projections from the IL synapse onto GABAergic PH neurons, likely contributing to the inhibitory tone within the nucleus (Myers et al., 2016). *In vivo* pharmacological experiments indicate that the PH is under tonic GABAergic inhibition, as microinjection of a GABA-A antagonist increases physiological stress responding in males (DiMicco et al., 1986; Myers et al., 2016).

Indeed, activation of the IL-PH circuit has divergent effects on stress responses to acute restraint in male and female rats, with stress-buffering effects in males and stress-enhancing effects in females in terms of corticosterone, blood glucose, heart rate, and motivated behaviors (Schaeuble et al., 2024). The neural mechanisms underlying this sexual divergence are unclear. In previously unstressed males and females, social, novel, and stress stimuli evoke similar population activity within IL neurons (Wallace and Myers, 2023), suggesting output circuitry may account for sex-specific regulation. Therefore, we hypothesize that sexual divergence in cortical-hypothalamic stress integration lies within circuit organization and signaling.

Here, we use slice electrophysiology and anatomical tracing to delineate the mPFC-PH circuit in male and female rats. Spontaneous PH firing properties were determined in sex- and cell-type manner using a glutamic acid decarboxylase (GAD)-Cre transgenic rat and Cre-dependent colorswitch virus. Channelrhodopsin (ChR2)-assisted circuit mapping (CRACM) was used to identify functional connections between IL and cell-type-defined PH neurons. Additionally, this circuit was mapped both anterogradely and retrogradely, with identification of stress-sensitive, hypothalamic-projecting neurons within mPFC subregions. Together, these data provide sex-specific functional organization of cortical-posterior hypothalamic circuity that modulates behavioral and physiological integration.

## Methods

### Animals

Male and female Long-Evans and Sprague-Dawley rats were used due to the availability of transgenic rats on a Long-Evans background and prior PH literature primarily being conducted in Sprague-Dawley rats (Abrahamson and Moore, 2001; Myers et al., 2016; Nyhuis et al., 2016; Schaeuble et al., 2024). For all experiments, rats were housed in shoebox cages with plastic tube enrichment. Sprague-Dawley rats were housed separately from Long-Evans rats to minimize stress; additionally male and female rats were housed in separate rooms. All rats were housed in temperature- and humidity-controlled rooms with a 12-hour light-dark cycle (lights on at 07:00 h, off at 19:00 h) with *ad libitum* food and water. All procedures and protocols are approved by the Colorado State University Institutional Animal Care and Use Committee (protocols 4994 and 4748) and complied with the National Institutes of Health Guidelines for the Care and Use of Laboratory Animals. Signs of poor health and/or weight loss ≥ 20% of pre-surgical weight were *a priori* exclusion criteria. Care was taken to minimize discomfort.

### Experiment 1: Intrinsic and Synaptic Properties of PH Neurons

#### Electrophysiology

##### PH Slice Preparation

Brain slices were prepared from Long-Evans (N = 5 male, 5 female, p112-164) and Sprague-Dawley (N = 5 male, 6 female, p72-166) rats. Spontaneous firing properties and synaptic inputs were examined in both Sprague-Dawley and Long-Evans rats to test for potential strain differences. Action potential features and synaptic inputs were measured in equal-sized groups (N = 10-11) of half male and half female rats in each strain, there were no differences that were statistically significant, so both strains were pooled to analyze potential sex differences.

Animals were deeply anesthetized with isoflurane, brains were removed and immediately submerged in ice-cold sucrose-based cutting solution containing the following (in mM): 83 NaCl, 2.5 KCl, 1 NaH_2_PO_4_, 26.2 NaHCO_3_, 22 D-Glucose, 72 sucrose, 0.5 CaCl_2_, 3.3 MgCl_2_. A block containing the PH was mounted on a vibrating microtome (VF-510-0Z Compresstome, Precisionary Instruments, Greenville, NC) and tissue was sliced at 300 µm into ice-cold sucrose-based cutting solution. Slices were transferred into artificial cerebrospinal fluid (aCSF) containing the following (in mM): 123 NaCl, 3.5 KCl, 1.25 NaH_2_PO_4_, 26 NaHCO_3_, 1 MgCl_2_, 10 D-glucose and 1.5 CaCl_2_. Slices were oxygenated with 95% O_2_ and 5% CO_2_ and incubated at 34°C for 30 minutes, then moved to room temperature.

##### Whole-Cell Patch Clamp Recordings

Slices were transferred to a recording chamber on an Olympus BX51WI upright microscope (Olympus, Tokyo, Japan) and superfused with aCSF maintained between 32°C and 34°C with an in-line heater (Warner Instruments, Holliston, MA). Patch-clamp electrodes were pulled from borosilicate glass (diameter, 1.5 mm; Sutter, Novato, CA) on a two-stage puller (Model PC-100, Narishige, Tokyo, Japan) and contained K-gluconate internal solution containing (in mM): 130 K-gluconate, 2 Na-gluconate, 6 NaCl, 2 MgCl_2_, 0.1 CaCl_2_, 1 EGTA, 4 MgATP, 0.3 trisGTP, 14 Tris-creatine phosphate, 10 sucrose, and 10 HEPES. Current clamp and voltage clamp recordings were made in whole-cell configuration using an amplifier (model Axopatch 200B, Molecular Devices, San Jose, CA) and Digidata 1440A digitizer (Molecular Devices). Recordings were excluded from analysis if access resistance was > 25 MΩ or leak current > 150 pA.

To quantify spontaneous inputs, cells were voltage clamped at -70 mV to record spontaneous excitatory postsynaptic currents (sEPSCs) and -5 mV to record spontaneous inhibitory postsynaptic currents (sIPSCs). 10 mV liquid junction potential was corrected on the amplifier and 5 mV correction was adjusted post-hoc. Intrinsic properties were calculated by P-clamp software (Molecular Devices) upon breaking in, action potential properties were collected in current clamp at I=0. Rheobase, sag, and FI curve data were collected in current clamp with current injections. To calculate rheobase, the cell was held at I=0, a -100 pA current injected for 100 ms, current then ramped from -100 pA to +100 pA at a constant rate of 0.8 pA/ms over 250 ms before returning to I=0. Rheobase was calculated as the current injection at which the first action potential reached threshold, averaged over 10 sweeps per cell. For the FI curve, current steps from -100 pA to +100 pA were injected in 10 pA increments from I=0, if depolarization block occurred at the end of the protocol, steps were removed from analysis. Sag was calculated from the most hyperpolarized (-100 pA) step in the FI protocol. Spontaneous action potential firing rate was calculated from I=0, action potential shape was used to make phase-plane plot and calculate maximum dV/dt, minimum dV/dt, as well as action potential halfwidth and amplitude from resting membrane potential.

##### Drugs

To test for GABA tone, the GABA_A_ receptor antagonist SR 95531 hydrobromide (gabazine; 10 µM; Tocris, Bristol, UK) was added to aCSF and superfused over the slice.

#### Estrous Cycle Cytology

Vaginal cytology was examined to approximate the estrous cycle stage. A cotton swab dampened with deionized water was used to collect cells from the vaginal canal and roll them onto a glass slide immediately after sacrifice. Dried slides were stained with H&E then viewed under a 10x objective light microscope by a minimum of two blind observers and were categorized as proestrus, estrus, metestrus, or diestrus (Cora et al., 2015; Schaeuble et al., 2023). Phases were grouped into those in stages of higher circulating hormones (proestrus & estrus, P/E) versus those in stages of lower circulating hormones (metestrus & diestrus, M/D; Egan et al., 2019).

Experimental design did not control for estrous phase, data were analyzed for cycle post hoc if N ≥ 3 per cycle. Data were split by estrous cycle if there was a significant effect of cycle. If there were no cycle effects, data were collapsed and cycle phase was highlighted by different symbols (empty upside-down triangle, M/D; filled in upside-down triangle, P/E).

#### Ovarian Hormone Receptor RNA

A subset of brains were prepared for transcriptional analysis using the Nanostring nCounter (Bruker Spatial Research, Bothell, WA) gene expression profiling system. Female Sprague-Dawley rats were deeply anesthetized with isofluorane, brains were collected and flash frozen with 2-methylbutane, then mounted in a cryostat where the PH was punched with a 700-μm tissue punch. Samples were prepared for transcriptional analysis via manufacturer’s guidelines at the University of Arizona Core Facilities (Tuscon, Arizona).

### Experiment 2: Cell-Type Specific Firing Properties

#### Animals

Long-Evans GAD-Cre transgenic rats were used to investigate cell-type specific firing properties (N= 5 male, 4 female; p77-110). This transgenic line was made by the Transgenic Rat Project at the National Institute of Drug Abuse (NIDA). GAD-Cre male rats (LE-Tg(GAD1-iCre)3Ottc) were obtained from the Rat Resource and Research Center (RRRC) at the University of Missouri. Long-Evans wild-type female rats were obtained from Charles River and bred in-house with Cre-positive males according to NIH recommendations. Rats were group-housed prior to surgery and single-housed following surgery. The GAD-Cre rat has been previously used and validated(Sharpe et al., 2017; Gibson et al., 2018; Luo et al., 2020; Prasad et al., 2020; Farrell et al., 2022), with 80% ± 6% of GAD1 cells co-expressing Cre in the LH (Sharpe et al., 2017). The identity of the Cre-, GFP-expressing cells have not been verified within the PH; therefore, we conducted histological analysis to determine the phenotype of these cells.

#### Stereotaxic Surgery

Rats were anesthetized with isoflurane (1-5%) followed by analgesic administration (0.6 mg/kg buprenorphine-SR, subcutaneous). AAV Nuc-flox(mCherry)-eGFP virus (Colorswitch; Back et al., 2019) was obtained from Addgene (Watertown, MA). Rats received bilateral microinjections of colorswitch virus (0.75 μL) directed at the PH with coordinates (mm from bregma): 4.0 caudal, ±0.5 lateral, 7.5 ventral from dura (Schaeuble et al., 2023). All microinjections were carried out with a 25-gauge, 2-μL microsyringe (Hamilton, Reno, NV) using a microinjection unit (Kopf, Tujunga, CA) at a rate of 5 min/μL. The needle was left in place for 5 min before and after injections to reduce tissue damage and allow diffusion. Skin was closed with wound clips that were removed 10-14 days following surgery.

#### Electrophysiology

Electrophysiological studies were conducted ∼5 weeks after viral injection. Patch clamp techniques were identical to intrinsic properties methods with the addition of identifying cell types prior to patching. GFP- or mCherry-expressing nuclei were identified with an X-cite Xylis laser (Excelitas Technologies, Pittsburg, PA).

#### RNAscope Tissue Preparation

A subset of brains was prepared for RNAscope histological analysis as follows: rats were given an overdose of sodium pentobarbital and transcardially perfused with 0.9% saline followed by 4% paraformaldehyde (PFA) in 0.1 M phosphate buffer solution (PBS). Brains were removed and post-fixed overnight at room temperature. The following day, brains were transferred into RNase-free, DEPC treated sucrose and stored at 4°C until slicing. Brains were flash frozen, sliced using RNase-free technique on a freezing microtome at 20 µm into RNase-free cryoprotectant (DEPC-treated RNase-free PBS, sucrose, PVP-40, ethylene glycol) and stored at -20°C.

#### Histology

##### RNAscope

The phenotype of GFP-expressing cells was determined using RNAscope. The presence of *vGluT2* transcript and *GAD2* transcript was quantified to identify glutamatergic and GABAergic neurons, respectively. RNAscope was conducted according to manufacturer’s instructions (ACD Bio; RNAscope Multiple Fluorescent V2 Assay) with minor modifications. Briefly, floating slices were washed in RNase-free PBS, transferred to hydrogen peroxide, and washed in RNase-free PBS prior to mounting on Superfrost Plus slides. Slices were allowed to dry on slides, then dehydrated in graded ethanol and dried at room temperature overnight. The following day, protease treatment, probe hybridization, preamplification, and amplification steps were performed according to manufacturer’s guidelines. Probes were fluorescently labeled with Opal 690 (1:300; Akoya Biosciences). Slides were coverslipped with Prolong Gold antifade (Thermo Fisher Scientific; Eugene, OR) and fluorescent images were captured on a Zeiss Axio Imager Z2 microscope and imaged at 63X with an oil-immersion objective. Microscopy images were exported from Zen and brightness was adjusted in Adobe Photoshop; adjustments were made to all channels equally. GFP co-localization with *vGluT2* or *GAD2* transcript was quantified by two experimenters blinded to condition.

##### Biocytin Immunohistochemistry

Recording pipettes contained internal solution with 2% biocytin (2 mg/mL; Sigma-Aldrich, St. Louis, MO) that passively diffused into neurons during recording for post hoc visualization. After recording, slices were post-fixed in 4% paraformaldehyde for 24-48 h at 4°C, then stored in 1% PBS at 4°C until biocytin immunohistochemistry. Slices containing biocytin-filled cells were processed according to previously published protocols (Davis et al., 2003). Briefly, slices were removed from PBS and incubated in electrophoretic tissue clearing solution (Logos Biosystems; catalog #C13001) for at least 2 h then washed in 1x phosphate buffered saline (PBS). Slices were then incubated in Streptavidin (Cy5, 1:1000 in 1% PBST; Vector Laboratories) for 2 h while shaking at room temperature, washed in 1x PBS (3 x 20 min) then coverslipped with polyvinyl mounting medium. Fluorescent images were captured on a Zeiss Axio Imager Z2 microscope and imaged at 10X. Images were exported from Zen and brightness was adjusted in Adobe Photoshop; adjustments were made to all channels equally.

##### Immunohistochemistry

To further determine the selectivity of the colorswitch virus, virally transfected tissue from unpatched electrophysiology slices was stained for microglial marker ionized calcium-binding adapter molecule 1 (Iba1), astrocytic marker glial fibrillary acidic protein (GFAP), or the rate limiting enzyme for dopamine production, tyrosine hydroxylase (TH) protein expression. PH slices that were sliced at 300 µm but not used for electrophysiology experiments were transferred to 4% PFA at the end of the recording day and post-fixed. Given the thickness of the tissue, previously published protocols were modified to increase antibody penetrance. Tissue was washed in 1X PBS for 10 min then incubated in 0.05 M PBS and 0.1 M glycine for 30 min before being washed in 1X PBS (3x5 min). Slices were then incubated in 0.5% sodium borohydride for 15 min before more washes in 1X PBS (3x5 min), then blocked in 5% normal goat serum with Triton X-100 for at least 30 min. Tissue was then incubated in primary antibody for 2 nights at 4°C with gentle agitation followed by washes in1X PBS (4 x 15 min), 1% normal goat serum, and 0.02% Triton X-100. The tissue was incubated in secondary antibody for 2h at 4°C, followed by 1X PBS washes (4x15 min) and 0.02% Triton X-100 before DAPI staining (10 min). Finally, tissue was washed in PBS and 0.02% Triton X-100 (4x15 min), mounted and coverslipped with polyvinyl alcohol mounting medium. Fluorescent images were captured on a Zeiss Axio Imager Z2 microscope and imaged at 40X. Images were exported from Zen and brightness was adjusted in Adobe Photoshop; adjustments were made to all channels equally.

GFAP was visualized with rabbit anti-GFAP primary antibody (1:1000; Dako, Carpinteria, CA) and Cy5 donkey anti-rabbit secondary (1:500; Jackson Immunoresearch, West Grove, PA). Iba-1 was labeled with rabbit anti-Iba-1 primary antibody (1:1000, Fujifilm Wako, Richmond, VA; Cat. # 019-19741) and Cy5 donkey anti-rabbit secondary (1:500; Jackson Immunoresearch,

West Grove, PA). TH was labeled with mouse anti-TH primary (1:5000; Sigma Aldrich, St. Louis, MO) and Cy5 Donkey anti-mouse secondary (1:500, Jackson Immuno Research, West Grove, PA).

GABA was immunolabeled as previously described (Schaeuble et al., 2024) to examine co-expression with mCherry-expressing nuclei in 20 µm thick tissue expressing the colorswitch virus. Briefly, coronal PH sections were removed from cryoprotectant and rinsed at room temperature in 1X PBS (5 x 5 min). Sections were then submerged into blocking solution (1X PBS, 6% bovine serum albumin, 8% normal goat serum, 2% normal donkey serum, and 0.4% Triton X-100) for 4h. Then, the tissue was incubated in rabbit anti-GABA primary antibody (Sigma, Saint Louis, Missouri; MFCD00162297) (1:250 in blocking solution) for 60 h at 4°C. After, the tissue was washed in PBS (5 x 5 min) and incubated in biotinylated secondary antibody goat anti-rabbit (Vector Laboratories; BA-100) (1:500 in PBS) for 2 h. Another PBS (5 x 5) wash was conducted before the tissue was incubated in Avidin-Biotin Complex (Vector Laboratories, Newark, California, Vectastain ABC Kit PK-4000) (1:500 in 1X PBS) for 1 h. The tissue was washed in PBS again (5 x 5) before incubation in Cy3-Streptavidin (Jackson Immuno Research; 016-160-084) (1:500 in PBS) for 1 h. Finally, the tissue was washed in PBS (5 x 5), mounted and coverslipped with polyvinyl alcohol mounting medium.

NeuN immunohistochemistry was conducted after RNAscope to identify neurons. After the final HRP blocker, slices were washed in PBS (5 x 5 min) on the shaker at room temperature, then incubated in blocking solution (1X PBS, 0.1% BSA, 0.2% Triton X-100) for 1 h. Slices were subsequently incubated in primary antibody (rabbit polyclonal anti-NeuN; Millipore, Damstadt, Germany; 1:2000) overnight. The following day, tissue was washed in 1X PBS (5 x 5 min) then incubated in secondary antibody (Cy3 donkey anti-rabbit; Jackson ImmunoResearch, Chester County, PA; 1:500) for 1 h, followed by another round of 1X PBS washes (5 x 5 min). Tissue was then coverslippled with polyvinyl alcohol mounting medium and imaged with a 40X objective.

### Experiment 3: Circuit-Specific Manipulations

#### Animals

Male and female transgenic GAD-Cre Long-Evans rats were used for all circuit-based electrophysiological experiments (N = 8 males; 7 females, p95-163).

#### Stereotaxic Surgery

Colorswitch virus was targeted to the PH, as described above. ChR2, the light sensitive non-selective cation channel, was targeted at the IL with coordinates (mm from bregma): 2.7 A/P, ± 0.6 lateral, 4.0 ventral from dura with a volume of 1.25 µL in males and 2.45 anterior, ± 0.5 lateral, 4.0 ventral from dura with a volume 1 µL in females (Schaeuble et al., 2024). This volume was used to facilitate diffusion laterally across cortical layers and horizontally through the rostral-caudal aspects of the IL. However, the volume allowed diffusion dorsally into the ventral aspects of the prelimbic cortex (PL), as well as some spread ventrally into the dorsal tenia tecta (TT), so the injection site was deemed mPFC. Injection site within the mPFC was confirmed post-hoc on Zeiss Axio Imager Z2 microscope, all injections successfully targeted the mPFC. The Swanson brain atlas (Swanson, 2018) was used to create viral injection brain maps.

ChR2 under the synapsin promoter (AAV5-hSyn-ChR2) was obtained from the University of North Carolina Vector Core (Chapel Hill, NC). Electrophysiological experiments were conducted 9-13 weeks after viral injection to allow time for ChR2 trafficking in this long-range projection. One electrophysiology experiment was conducted after 4 weeks of viral incubation to test incubation time, while the optically-evoked excitatory postsynaptic currents (oEPSCs) did not show robust differences, the oEPSCs were more reliable after the longer time course, so a 9-13 week viral incubation time was used for the remainder of the experiments. This long incubation time is consistent with previously published work on similar circuits and consistent with long-range transport required for such a projection (Warden et al., 2012).

#### Electrophysiology

For ChR2 oEPSCs, recording electrodes were filled with Cs-MeSO_4_ internal solution containing the following (in mM): 120 MeSO_4_, 3 NaCl, 2 MgCl_2_, 1 EGTA, 10 HEPES, 4 MgATP, 0.3 TrisGTP, 14 Tris-creatine phosphate, 12 sucrose, 1.2 QX314, 4 TEA-Cl. Slices were incubated in and superfused with aCSF containing the following (in mM): 123 NaCl, 3.5 KCl, 1.25 NaH_2_PO_4_, 26 NaHCO_3_, 1 MgCl_2_, 10 D-glucose and 2 CaCl_2_. The CaCl_2_ concentration was adjusted slightly to increase the probability of release. oEPSCs were stimulated with 470nm light (Thor Labs LEDD1B; Newton, NJ). GFP- and mCherry-expressing cells were visualized with the X-Cite laser, as described above. Cells were passively filled with 2% biocytin throughout the course of the experiment for post-hoc phenotype confirmation, as described above.

##### Pharmacology

Identity of the mPFC inputs was determined pharmacologically. Glutamatergic signaling was blocked with a bath application of the AMPA receptor antagonist 6,7-dinitroquinoxaline-2,3-dione (DNQX; 10 µM; Tocris Biosciences, Bristol, UK) and the NMDA receptor antagonist 3-((+)-2-carboxypiperazin-4-yl)propyl-1-phosphonic acid (CPP; 10 µM; Tocris Biosciences). In a subset of experiments, GABAergic signaling was blocked with bath application of the GABA_A_ receptor antagonist SR 95531 hydrobromide (gabazine; 10 µM; Tocris Biosciences).

### Experiment 4: Organization of the mPFC-PH Circuit

#### Animals

Sprague-Dawley rats were used for retrograde (N = 5 male, 6 female) and anterograde (N = 4 male, 5 female) anatomical tracing studies. Animals arrived 6-8 weeks old, were allowed 1 week to acclimate prior to surgery, and euthanized with an overdose of sodium pentobarbital 6 weeks following viral injection.

#### Stereotaxic Surgery

##### Retrograde Tracing

Bilateral injections of a retrograde AAV were targeted to the PH (coordinates from Bregma in mm): 4.0 posterior, ± 0.6 lateral, 7.5 dorsal from dura; 0.75 µL bilaterally). The retrograde-transported virus used for viral tracing (pAAV-hSyn-Cre-P2A-dTomato; 107738-AAVrg) carrying a construct coding for Cre and dTomato was obtained from Addgene (Schaeuble et al., 2024).

##### Anterograde Tracing

Synaptotag virus was injected targeting the IL with coordinates (in mm from Bregma): 2.6 rostral,0.6 lateral, 4.2 ventral from dura; 50 nL bilaterally. Brains were collected 6 weeks after microinjection to allow time for construct expression. Synaptotag (mCh-GFP-syb2) was obtained from the Stanford Gene Vector and Virus Core (Palo Alto, CA) and carries a construct under the control of the human synapsin promoter (hSyn1) (Pace et al., 2020). The virus codes for mCherry expression in neuronal soma and axons and enhanced GFP conjugated to synaptobrevin-2, creating a distinction between soma / axons and presynaptic terminals. Fluorescent images were captured on a Zeiss Axio Imager Z2 microscope at 40x with an oil-immersion objective. Images were exported from Zen and brightness was adjusted in Adobe Photoshop; adjustments were made to all channels equally.

#### Histology

Transcardial perfusions were conducted and brains were removed 90 minutes after the initiation of an acute, 30-minute restraint stress, as previously described (Schaeuble et al., 2024). Brains were prepared as described above and sliced at 30 μm. Immunohistochemical staining for c-Fos protein expression was conducted as previously published (Myers et al., 2016; Schaeuble et al., 2024) to quantify co-localization of c-Fos and retrograde tracer. Floating sections containing forebrain were removed from cryoprotectant and washed in 1x PBS at room temperature (5x5min). Tissue sections were placed in blocking solution (1X PBS, 4% goat serum, 0.4% Triton-X-100, 0.2% bovine serum albumin) for 30 min. After blocking, the tissue was incubated in primary mouse-anti-Cre recombinase antibody (1:1000; Millipore) overnight at 4 degrees to boost signal relative to dTomato alone. The following day, the tissue sections were washed in 1x PBS (5x5min) then incubated in Cy3 goat anti-mouse secondary antibody (Jackson Immuno Research, West Grove Pennsylvania, 115-165-003, 1:800 in 1xPBS) for 1 hour. Next, sections were washed in 1x PBS 5 x 5 min and placed in blocking solution (1xPBS, 0.1% BSA+0.4% Triton-X100) for 1 hour. Once blocked, the tissue was incubated in polyclonal guinea pig anti-c-Fos primary antibody (1:1000; Synaptic Systems, Gottingen, Germany) overnight at 4 degrees. The next day, the tissue sections were washed in 1xPBS (5x5min) then incubated in Cy5 donkey-anti-guinea pig secondary antibody (1:1000; Synaptic Systems, Gottingen Germany) for 1 hour. Finally, the tissue was washed in 1x PBS, mounted and cover slipped with polyvinyl mounting medium. Fluorescent images were captured on a Zeiss Axio Imager Z2 microscope at 63x with an oil-immersion objective. The number of retrograde labeled cells within mPFC subregions and co-localization with cFos protein were quantified by two experimenters blinded to conditions. Images were exported from Zen and brightness was adjusted in Adobe Photoshop; adjustments were made to all channels equally.

To confirm the presynaptic nature of GFP-synaptobrevin-2 expression and the glutamatergic neurochemistry of mPFC inputs to the PH, SynaptoTag tissue was immunolabeled for vesicular glutamate transporter 1 (vGluT1) (Myers et al., 2017). Coronal forebrain sections were rinsed in 1x PBS (5x5min) to remove cryoprotectant solution. Tissue sections were then incubated for 1 hour at room temperature in blocking solution (1x PBS, 0.1% bovine serum albumin, 0.2% TritonX-100). Following blocking, sections were incubated in rabbit anti-vGluT1 primary antibody (1:1000 in blocking solution: Synaptic Systems, Göttingen, Germany; 135-330) overnight at 4 degrees. The following day, the tissue was washed in 1x PBS (5x5min) then incubated in Cy5 conjugated donkey-anti-rabbit secondary antibody (1:500 in 1x PBS: Jackson Immuno-Research, West Grove, PA) for 30 minutes at room temperature. After incubation, the tissue underwent a final round of 1x PBS washes (5x5min) and was mounted then coverslipped with polyvinyl alcohol mounting medium and imaged on a Zeiss Axio Imager Z2 microscope at 63x with an oil-immersion objective.

### General Methods

#### Data Analysis

All data are presented as mean ± standard error of the mean. Data were analyzed in Prism 10.4.1 (GraphPad, San Diego, CA), with statistical significance set at *p* < 0.05 for rejection of the null hypotheses. Statistically significant results are highlighted in figures, statistical tests resulting in 0.05 < p < 0.06 are highlighted in the results section. Parametric statistical tests were conducted via unpaired *t* test, one way ANOVA, 2-way ANOVA, or 2-way repeated measures ANOVA. Post-hoc analyses were run when ANOVA results indicated a main effect or interaction between factors, specific post-hoc analyses are detailed in the results section. Binary data were analyzed with a Fisher’s exact test. When analysis revealed a significant result, pairwise Fisher’s exact test post-hoc tests were run.

## Results

### Intrinsic and Synaptic Properties of PH Neurons

As there are no published reports of intrinsic cell physiology within the PH to our knowledge, we examined spontaneous action potential firing, excitability, and excitatory / inhibitory balance to provide a foundation for cell-type specific analysis and examination of circuit signaling. Spontaneous firing properties were examined in Sprague-Dawley and Long-Evans rats to test for strain differences, as previous literature examining PH stress regulation has been conducted in Sprague-Dawley rats (Myers et al., 2016; Nyhuis et al., 2016; Schaeuble et al., 2024) but GAD-Cre transgenic animals are on a Long-Evans background (Sharpe et al., 2017). Action potential features and synaptic inputs were measured in equal-sized (N = 10-11) groups of half male and half female rats in each strain. There were no statistically significant differences, so both strains were pooled. Subsequent electrophysiological experiments used Long-Evans for transgenic availability. All analyses compared male to female, then compared female in metestrus/diestrus (m/d) versus females in proestrus/estrus (p/e). In the case of significant estrous effects, they were graphed separately. If no estrous effect was observed, the female data remained pooled with individual data points coded for m/d vs p/e. No estrous phase x sex comparisons were carried out.

PH neurons that fire at rest had a resting membrane potential of -61.66 ± 0.77 mV and were significantly more depolarized than PH neurons that were quiet at rest, which had a resting membrane potential of -73.14 ± 1.96 mV (**Fig. 1A**; unpaired *t* test, *t*(104)=4.089, *p* < 0.0001). However, there were no differences between the resting membrane potential of PH neurons from male and female rats (**Fig. 1B**; male = -63.93 ± 1.09, female = -62.68 ± 1.22; unpaired *t* test, *t*(104) = 0.7636, *p* = 0.4468). When females were split by estrous cycle, there was a trend toward a more depolarized resting membrane potential in females with lower levels of circulating hormones (Female M/D: -59.06 mV, Female P/E: -64.20 mV; unpaired *t* test, t(52) = 1.981, *p* = 0.0529). Consistent with a similar resting membrane potential between the sexes, a similar proportion of PH neurons fired at rest (male 83%, female 85%; χ^2^ (1, n = 106) = 0.1223, *p* = 0.7266; **Fig. 1C**). While spontaneous actional potential firing frequency was not dependent on sex (unpaired *t* test, *t*(87) = 1.310, *p* = 0.1937), within females, firing frequency was dependent on estrous cycle. The spontaneous rate of action potentials in M/D females was 25.40 ± 5.13 Hz while the spontaneous firing rate of neurons from females in P/E was 13.66 ± 2.35 Hz (unpaired *t* test, *t*(43) = 2.283), while PH neurons from male rats fire at 13.39 ± 1.671 Hz (**Fig. 1D**). Additionally, there was a significant difference in the capacitance of PH neurons from male and female rats (male = 35.16 ± 1.304 pF, female = 29.64 ± 1.352 pF; unpaired *t* test, *t*(104) = 2.934; **Fig. 1E**).

**Figure 1.**
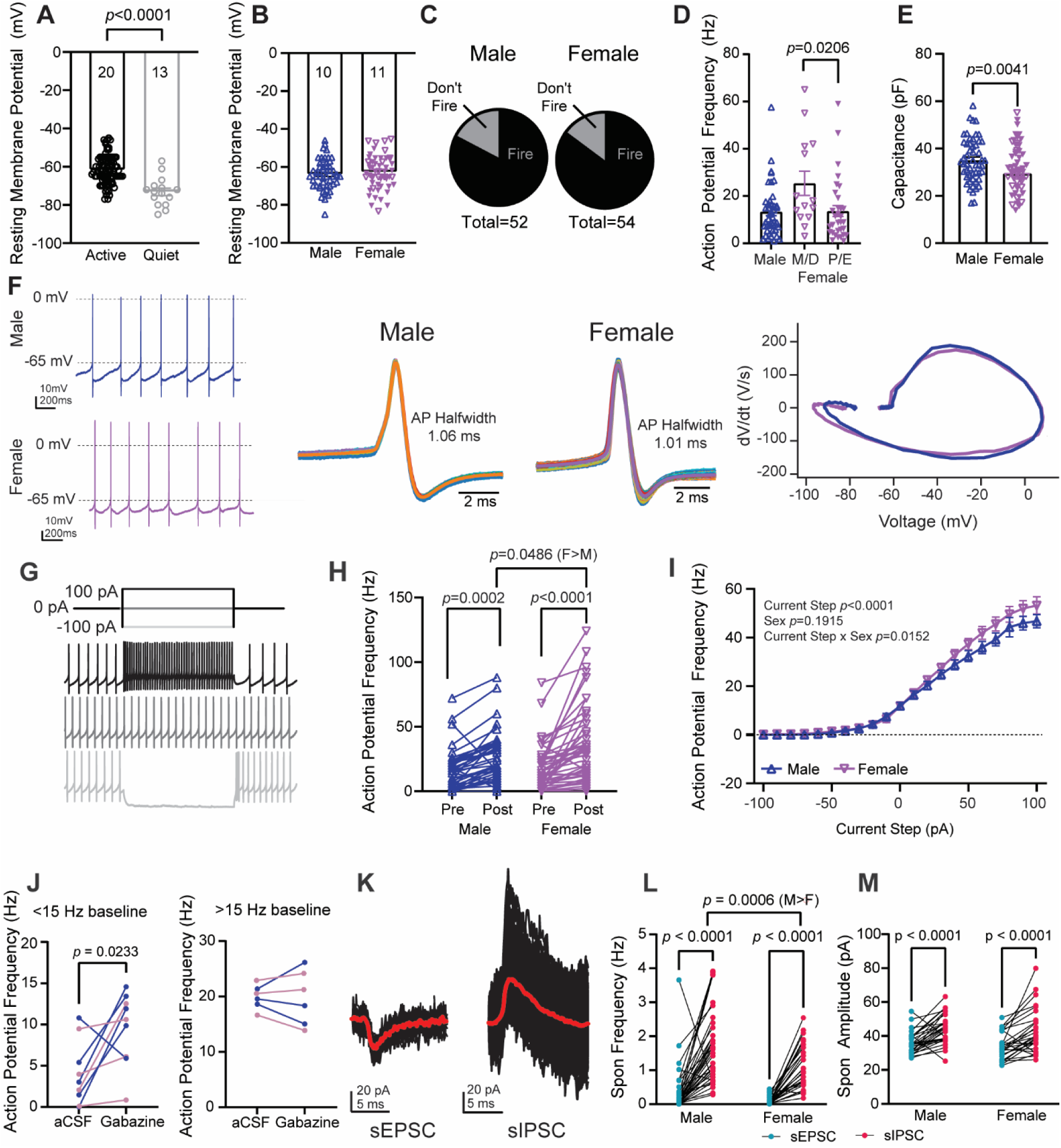
Firing properties of neurons within the PH of male and female rats. **A.** Resting membrane potential of neurons that are active or quiet at rest with sex collapsed; **B.** Resting membrane potential of neurons from male and female rats; **C.** Proportion of cells that fire at rest; **D.** Action potential frequency at I=0; **E.** Capacitance; **F.** Representative action potential firing at I=0, action potential waveform, and dV/dt plot, P/E female; **G.** Representative FI curve showing -100 pA, 0 pA, and +100 pA current injections; **H.** Action potential frequency before (pre) and after (post) -100 pA current injection, N=10 male, 10 female; n= 44 cells from male, 48 cells from female; I. FI Curve, N=10 male, 10 female; n=44 cells from male, 48 cells from female; **J.** Action potential frequency before and after superfusion of gabazine (10 μM) separated by whether the cell fires below (<15 Hz; N = 8 cells from 5 animals) or above (>15 Hz; 4 cells from 3 animals) the average firing frequency of PH neurons; **K.** Overlay of sEPSC and sIPSC events from one cell, average waveform in red; **L.** Spontaneous event frequency, lines connect sESPC and sIPSC from the same cell (N = male: 38 cells from 9 animals; female: 32 cells from 8 animals); **M.** Spontaneous event amplitude, lines connect sEPSC and sIPSC from the same cell (N = male: 31 cells from 9 animals; female: 27 cells from 8 animals). *P*-values < 0.05 are shown on graphs. If there was a significant difference between estrous cycles, the female group is split by estrous.

Action potentials were converted to phase-plane plots using dV/dt analysis to analyze characteristics of the action potential shape (**Fig. 1F**). All components of the action potential (halfwidth, height, max dV/dt, min dV/dt) were not significantly different between PH neurons from male and female rats (**Table 1**). Neural excitability was analyzed two ways: hyperpolarization-induced action potential firing and FI curve. Hyperpolarization-induced firing was measured by quantifying action potential frequency 0.5s prior to (pre) and immediately following (post) a 1s, -100pA current injection. In PH neurons from both males and females, there was a significant increase in action potential frequency after the step (post) compared to before (pre; **Fig. 1H**). Interestingly, action potential frequency immediately following the hyperpolarizing step was significantly higher in females compared to males (male *n* = 44 cells, female *n* = 48 cells; male post-step frequency, 25.45 ± 2.96 Hz; female post-step frequency, 34.46 ± 4.35 Hz; 2-way ANOVA main effect of pre v post: F_(1, 90)_ = 59.56, *p* < 0.0001, interaction: F_(1, 90)_ = 4.056, *p* = 0.0470, Fishers LSD post hoc: female pre-vs-post *p* < 0.0001, male pre-vs-post *p* = 0.0002, post male-vs-female *p* = 0.0486). There was also a trend towards a cycle effect with higher pre-step firing frequency in M/D females compared to P/E females (2-way repeated measures ANOVA, main effect of pre-v-post F(1,46) = 22.70, *p* < 0.0001, Fisher’s LSD test, pre-female M/D vs. female P/E, *p* = 0.0501). This trend is consistent with the baseline action potential frequency. Action potential frequency throughout the FI curve current injections revealed a significant current by sex interaction (2-way repeated measures ANOVA; current x sex interaction F(20, 1730) = 1.812, *p* = 0.0152; **Fig. 1I**). Sag, a measure of HCN channel activity, was analyzed during a -100 pA current step. Only 10 of 44 cells from male and 13 of 49 cells from female PH exhibited sag (> 5mV change from the start of the step to steady state during the step). Of the cells that did exhibit sag, there was no difference in the magnitude of the sag between cells from male and female rats (**Table 1**). There was also no difference in rheobase between PH cells from male and female rats (**Table 1**). Together, these data indicate that PH spontaneous firing and membrane properties can depend on biological sex or estrous cycle, but many properties are not dependent on either. Additionally, neurons from the female PH exhibit increased excitability compared to those from the male PH.

**Table 1:** Spontaneous membrane and action potential properties of unlabeled PH cells, sex analysis with strains pooled (N rats = 10 male, 11 female).

| Parameter | Male | Female | p-value | n cells |
| --- | --- | --- | --- | --- |
| *Intrinsic Properties* |  |  |  |  |
| Membrane Resistance (MΩ) | 694.5 ± 75.29 | 682.6 ± 60.25 | 0.9011 | 48, 53 |
| *Action Potential Properties* |  |  |  |  |
| Maximum dV/dt (V/s) | 147.0 ± 8.552 | 142.8 ± 8.130 | 0.7241 | 43, 46 |
| Minimum dV/dt (V/s) | -127.7 ± 5.949 | -121.3 ± 5.743 | 0.4448 | 43, 46 |
| Halfwidth (ms) | 1.028 ± 0.04614 | 1.151 ± 0.05623 | 0.0969 | 43, 46 |
| Amplitude (mV) from -70 mV | 87.90 ± 2.141 | 88.28 ± 2.333 | 0.9044 | 43, 46 |
| Sag (>5 pA; at -100 pA step) | 8.560 ± 0.9304 | 12.34 ± 1.960 | 0.1286 | 10, 13 |
| Rheobase from ramp (pA) | -5.231 ± 3.869 | -6.199 ± 3.604 | 0.8554 | 49, 47 |

### The PH is Under Tonic GABAergic Control

*In vivo* experiments indicate that the PH is under tonic GABAergic control. Pharmacologically relieving inhibitory tone reduces social behaviors and increases anxiety-like behaviors (Myers et al., 2016), as well as increasing heart rate and pressor responses in male rats (DiMicco and Abshire, 1987). To test the extent of this tone within the PH of male and female rats, action potential frequency was measured prior to and after bath application of the GABA-A receptor antagonist gabazine (10 μM; **Fig. 1J**). In neurons that fire below the mean firing frequency of 15 Hz, gabazine significantly increased firing frequency (paired *t* test, *t*(8) = 2.797, *p* = 0.0233). Interestingly, neurons that have a baseline firing rate over 15 Hz were not impacted by the addition of gabazine (paired *t* test, *t*(5) = 0.1066, *p* = 0.9192), indicating GABAergic tone onto a subset of PH neurons that may be dependent on spontaneous firing frequency. GABAergic tone was also tested by quantifying spontaneous inputs onto PH neurons. Spontaneous excitatory (sEPSC) and inhibitory (sIPSC) postsynaptic currents were measured in voltage clamp. To record sEPSCs and sIPSCs from the same neuron, the holding voltage was adjusted from -70 mV (sEPSCs) to -5 mV (sIPSCs). In both males and females, sIPSC frequency (**Fig. 1L**) and amplitude (**Fig. 1M**) was larger than sEPSCs (frequency: 2-way ANOVA, main effect of E/I: *F*_(1, 68)_ = 91.93, *p* < 0.0001, main effect of sex: *F*_(1, 68)_ = 7.554, *p* = 0.0077; amplitude: main effect of E/I: *F*_(1, 56)_ = 49.40, *p* < 0.0001). These data further support an inhibitory tone within the PH of both male and female rats. Interestingly, PH neurons from male rats have significantly elevated sIPSC frequency (Fisher’s LSD test: *p* = 0.0097), indicating a larger inhibitory tone in males compared to females.

### Ovarian Hormone Receptor Transcripts

Prior work investigating estrogen and progesterone receptor expression within the hypothalamus has focused predominantly on ovariectomized females and the anterior hypothalamus (Simerly et al., 1990; Shughrue and Merchenthaler, 2001). To investigate the relative abundance of estrogen and progesterone receptor transcripts in intact females that could impact spontaneous action potential frequency, we examined RNA in PH punches from freely-cycling female Sprague-Dawley rats. There was a significant difference in the expression of estrogen receptor α (*Esr1*), estrogen receptor β (*Esr2*), and progesterone receptor (*Prg*; one way ANOVA: F(2,21) = 31.12, *p* < 0.0001, **Fig. 2**). Tukey’s multiple comparisons test revealed that *Prg* RNA is significantly greater than both *Esr1* and *Esr2* transcripts, suggesting that *Prg* may be a predominant contributor to female PH estrous signaling.

**Figure 2.**
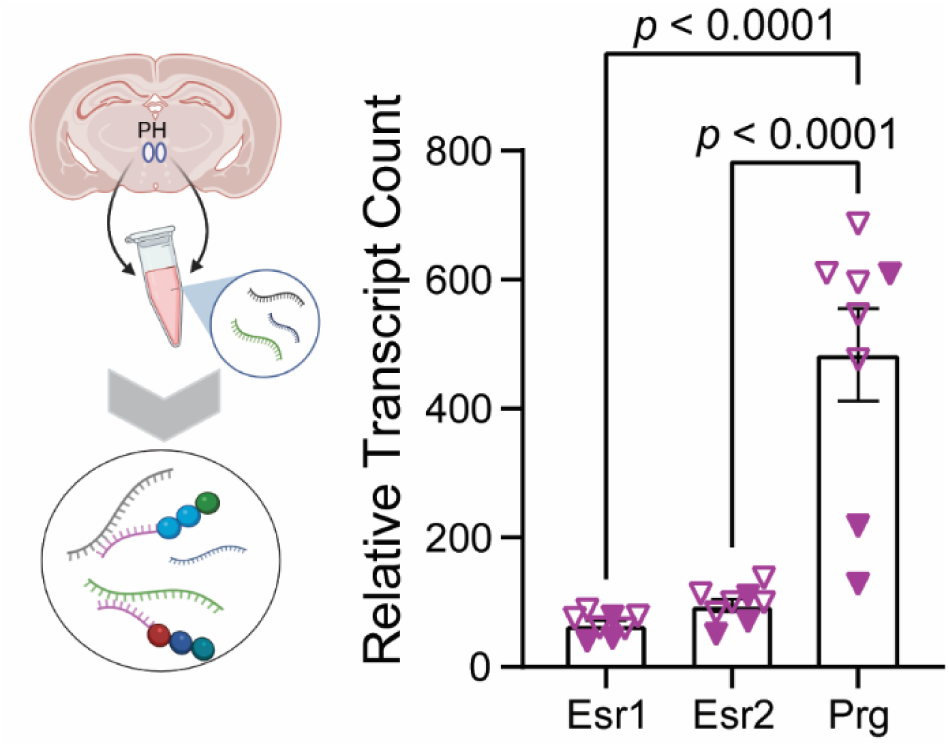
Estrogen receptors α, β and progesterone receptor RNA in the PH of freely-cycling females. Transcripts from PH punches were quantified for hormone receptor gene expression in freely-cycling females. Proportion of estrous phases at tissue collection: N = 5 M/D and N = 3 P/E. Esr1: ERα; Esr2: ERβ; Pgr: progesterone receptor.

### Cell-Type Specific Properties

The PH is composed of a heterogenous neuronal population, comprised primarily of glutamatergic and GABAergic neurons (Abrahamson and Moore, 2001). To investigate firing properties in a cell-type specific manner, a Cre-dependent colorswitch virus (AAV-Nuc-flox(mCherry)-eGFP) was microinjected into the PH of male and female GAD-Cre rats (Sharpe et al., 2017; Back et al., 2019). When expressed in the absence of Cre recombinase, this viral construct induces nuclear GFP expression. However, in the presence of Cre recombinase, the construct induces nuclear mCherry expression. This viral approach allowed for the visual identification of GAD+ and GAD- cells within the PH and cell-type specific electrophysiological recordings (**Fig. 3A, B**). The GAD-Cre rat has been extensively used and validated (Sharpe et al., 2017; Gibson et al., 2018; Luo et al., 2020; Prasad et al., 2020; Farrell et al., 2022) with 80% ± 6% co-expression of GAD1 and Cre in the LH (Sharpe et al., 2017). We confirmed the GABAergic nature of mCherry-expressing cells with GABA immunohistochemistry (**Fig. 3C**). Therefore, mCherry was used as a marker of GABAergic PH neurons. To identify the phenotype of GFP-expressing cells, RNAscope was used to quantify the proportion of GFP-expressing cells that express vGluT2 (s*lc17a6*) (El Mestikawy et al., 2011) or GAD65 (*gad2*) (Myers et al., 2014) transcripts, markers for glutamatergic and GABAergic neurons, respectively, within the PH (**Fig. 3D, E**). 77 ± 7% of GFP-expressing cells co-localized with vGluT2 transcript while 12 ± 3% co-localized with GAD2+ transcript (**Fig. 3F**), indicating that GFP cells are predominantly glutamatergic with a minor subset of GAD+ neurons, likely due to the absence of Cre or failed recombination of the colorswitch construct.

**Figure 3.**
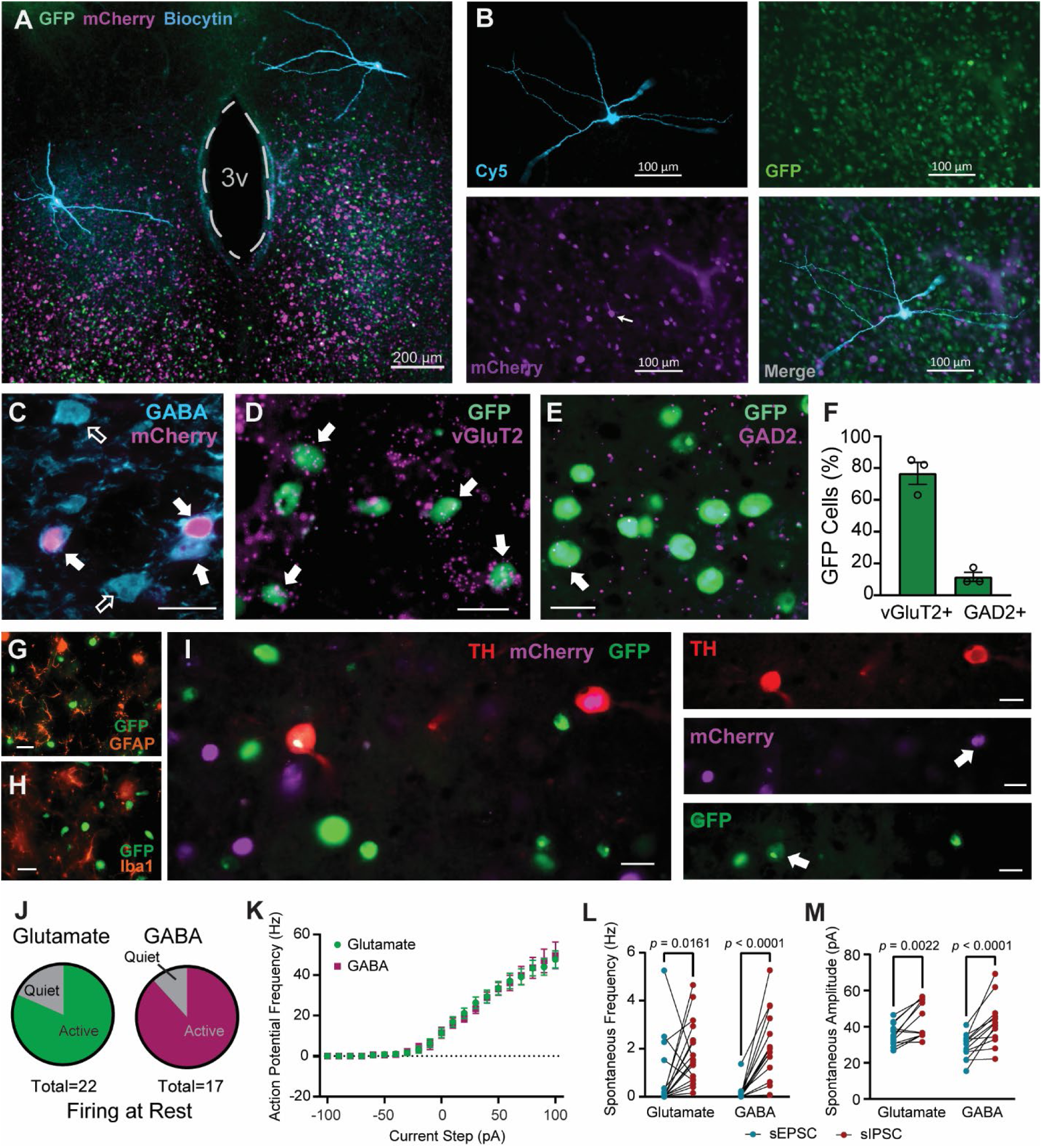
GAD-positive and GAD-negative neurons are predominantly GABAergic and glutamatergic, respectively, and not distinguishable by spontaneous properties. Cell-type specific firing properties within the PH. **A.** Biocytin-filled cells within the PH; **B.i-iv.** Biocytin-filled GAD+ neuron within the PH; **C.** Representative GABA IHC with mCherry neurons co-localizing (filled in arrows) with GABA immunoreactive cells (open arrows); **D.** vGluT2 RNAscope within the PH; **E.** GAD2 RNAscope within the PH; **F.** RNAscope quantification of colocalization of vGluT2 and GAD2 with GFP; **G.** GFAP immunohistochemistry within the PH; **H.** Iba1 immunohistochemistry within the PH; **I.** TH immunohistochemistry; **J.** Proportion of cells that fire at rest; **K.** FI curve (N: GFP = 21 cells from 9 animals; mCherry = 15 cells from 8 animals); **L.** Spontaneous excitatory and inhibitory postsynaptic current frequency, lines connect sEPSC and sIPSC from the same cell; **M.** Spontaneous excitatory and inhibitory postsynaptic current amplitude, lines connect sEPSC and sIPSC from the same cell. Unlabeled scale bar = 20 μm.

Immunohistochemistry for GFAP and Iba1 to identify astrocytes and microglia, respectively, was conducted to determine whether any proportion of GFP-expressing cells were glia (**Fig. 3G, H**). There was no co-localization between these glial markers and GFP-expressing nuclei, suggesting that the colorswitch construct was not expressed in glial cells within the PH. There is a sparce population of dopaminergic neurons within the PH (Abrahamson and Moore, 2001; Conceição Furber et al., 2021). To test whether these dopaminergic neurons comprise a selective subset of either mCherry- or GFP-expressing neurons, immunohistochemistry was conducted for TH, the rate-limiting enzyme for dopamine synthesis (**Fig. 3I**). TH immunoreactivity was co-expressed with both GFP and mCherry, suggesting that TH does not comprise a substantial portion of either cell population and is, instead, a subset of both GFP- and mCherry-expressing populations. These data validate the approach where GFP-expressing cells are predominantly glutamatergic and mCherry-expressing cells are predominantly GABAergic.

Interestingly, intrinsic properties and action potential properties were strikingly similar between glutamatergic and GABAergic cells within the PH (**Table 2**). Given the statistical similarities in spontaneous firing properties between males and females, sex was pooled to examine potential firing property differences between putative glutamatergic and GABAergic neurons (N rats = 4 males, 5 females). The majority of cells in both groups fire at rest (glutamate 82%, GABA 88%; χ^2^ (1, n = 39) = 0.3034, *p* = 0.5818; **Fig. 3J**) and have similar levels of excitability in response to current injection (2-way ANOVA, cell type main effect: F_(1, 34)_ = 0.005374; *p* = 0.9420; **Fig. 3K**). Both cell types receive more spontaneous inhibitory input than excitatory input, with higher frequency and larger amplitude sIPSCs compared to sEPSCs in both groups (Frequency: 2-way repeated measures ANOVA, main effect of excitatory / inhibitory F(1,30) = 25.64, *p* < 0.0001, Fishers LSD post hoc test: glutamate excitatory / inhibitory *p* = 0.0161, GABA excitatory / inhibitory *p* < 0.0001; Amplitude: 2-way repeated measures ANOVA, main effect of excitatory / inhibitory F(1,23) = 38.13, Fisher LSD post hoc test: glutamate excitatory / inhibitory *p* = 0.0022, GABA sEPSC v sIPSC *p* < 0.0001; **Fig. 3L, M**) further supporting an inhibitory tone within the PH, independent of postsynaptic cell type.

**Table 2:** Spontaneous membrane and action potential properties of defined cell types within the PH, sex pooled (N rats = 9 GFP, 8 mCherry). Membrane resistance and capacitance were calculated for all colorswitch cells (Fig. 2 and Fig. 3) therefore include a larger sample size (N rats = 27, 26).

| Parameter | GFP | mCherry | p-value | n cells |
| --- | --- | --- | --- | --- |
| *Intrinsic Properties* |  |  |  |  |
| Membrane Resistance (MΩ) | 973.4 ± 120.1 | 952.4 ± 123.0 | 0.9035 | 76, 64 |
| Capacitance (pF) | 36.72 ± 1.4 | 33.84 ± 1.44 | 0.1615 | 76, 64 |
| Resting Membrane Potential (mV) | -65.68 ± 1.5 | -65.32 ± 1.11 | 0.857 | 22, 17 |
| *Action Potential Properties* |  |  |  |  |
| Fire @ Rest Percentage | 82% | 88% | 0.679 | 22, 17 |
| Spontaneous Rate (spike / s) | 10.00 ± 2.35 | 8.05 ± 1.79 | 0.5341 | 22, 17 |
| Maximum dV/dt (V/s) | 168 ± 15.65 | 149.2 ± 15.07 | 0.3948 | 18, 16 |
| Minimum dV/dt (V/s) | -140.4 ± 10.63 | -128.3 ± 12.63 | 0.4673 | 18, 16 |
| Halfwidth (ms) | 0.99 ± 0.07 | 1.23 ± 0.14 | 0.1176 | 18, 16 |
| Amplitude (mV) from -70 mV | 100.2 ± 1.97 | 101.7 ± 2.47 | 0.6204 | 18, 16 |
| Rheobase from ramp (pA) | 1.60 ± 5.82 | 2.80 ± 4.54 | 0.8788 | 21, 16 |
| Sag (>5 pA; at -100 pA step) | 7.13 ± 1.11 | 12 ± 5.30 | 0.3399 | 4, 3 |

### Circuit-Specific Input to the PH

The PH receives dense input from the mPFC (Myers et al., 2016; Wood et al., 2019) and IL-PH activation can link cognitive and emotional processing to homeostasis and motivation, with different effects in male and female rats (Schaeuble et al., 2024). It is not clear, though, whether this information is conveyed through differential postsynaptic neuronal targeting via direct synaptic connection to distinct PH cell types. Previous work identified synaptic connections between IL inputs and PH GABA neurons (Myers et al., 2016). In addition, we combined histology with viral tracing from the mPFC and found that mPFC inputs also synapse onto glutamatergic PH neurons (**Fig. 4A**). To test the relative functional connectivity within this circuit, we used CRACM (CRACM; Petreanu et al., 2007; Atasoy et al., 2008; Garfield et al., 2015). ChR2 under the synapsin promoter was microinjected into the mPFC and colorswitch virus was microinjected into the PH of male and female GAD-Cre Long-Evans rats (**Fig. 4B**). Injection site and viral spread were restricted to the mPFC (**Fig. 4C**). Consistent with prior reports (Myers et al., 2017; Wood et al., 2019), we confirmed that the mPFC-PH projection is predominantly glutamatergic as ChR2 oEPSCs were blocked by glutamate receptor antagonists, DNQX and CPP (10 µM; **Fig. 4D**). Paired pulse ratio (PPR) was used to measure presynaptic release probability from mPFC inputs. No difference in PPR was observed between postsynaptic identity or sex (2-way ANOVA, main effect of sex F(1, 43) = 0.6383, *p* = 0.4287; main effect of cell type F(1,43) = 0.7265, *p* = 0.3987; **Fig. 4E**).

**Figure 4.**
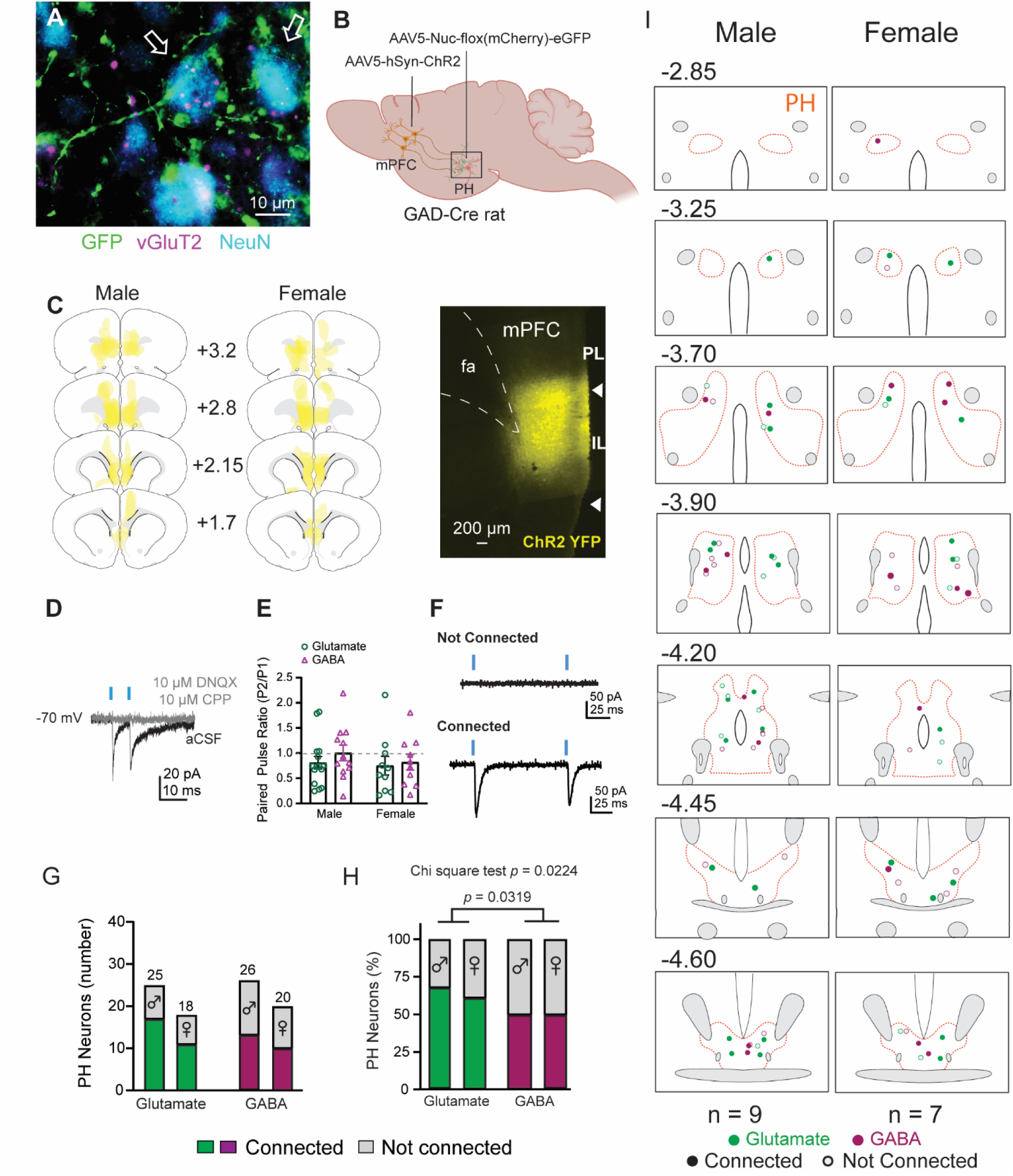
mPFC terminals target glutamatergic and GABAergic PH neurons. mPFC inputs onto identified vGluT2+ and vGluT2- neurons within the PH. **A.** Image of an IL terminal (GFP) impinging on vGluT2 mRNA-expressing neurons (magenta) in the PH (NeuN in blue); Schematic with viral constructs; **B.** Viral injection schematic; **C.** Injection map of mPFC YFP ChR2 injection placements and representative injection (2.8 mm from Bregma); **D.** Functional confirmation that mPFC inputs to the PH are glutamatergic, as they are blocked by bath application of NBQX (10 μM) and CPP (10 μM); **E.** Paired pulse ratio from cells that exhibited oEPSCs (N = Male GFP: 15 cells from 7 rats, Male mCherry: 13 cells from 6 rats, Female GFP: 10 cells from 7 rats, female mCherry: 10 cells from 6 rats); **F.** Representative traces from PH cells that do not receive input (not connected) and those that do receive input (connected) from mPFC afferents; **G.** Number of responder and non-responder PH neurons separated by sex and cell-type; **H.** Percentage of total cells that are responders or non-responders separated by sex and cell-type; **I.** Anatomical map of all recorded cells and whether they were connected or not connected to mPFC afferents. PL, prelimbic region of the prefrontal cortex; IL, infralimbic region of the prefrontal cortex.

Optical stimulation resulted in oEPSCs in a subset of neurons, indicative of a connection between mPFC terminals and postsynaptic neurons (**Fig 4F**). PH cells were then sorted into cells that are not connected and cells that are connected. Surprisingly, the proportion of glutamatergic and GABAergic cells targeted by mPFC inputs in males and females was not different (%; χ^2^ (3, n = 89) = 2.279, *p* = 0.5166; **Fig. 4G**). However, when data were normalized to assess the percentage of connectivity for each cell type, Chi-square test analysis revealed a significant difference in the percentage of connected neurons between the four groups (male v female, glutamate v GABA; %; χ^2^ (3, n = 89) = 9.592, *p* = 0.0224; **Fig. 4H**). Post hoc analysis revealed a higher proportion of connectivity onto glutamate neurons when sex is collapsed (χ^2^ (1, n = 89) = 4.604, *p* = 0.0319) and, specifically within males, a significantly larger connected proportion mPFC connectivity to glutamate neurons (χ^2^ (1, n = 51) = 0.6.697, *p* = 0.0097). To understand the rostral-caudal intranuclear distribution of PH glutamate and GABA cells and their responsivity to mPFC inputs, the location of patched cells was mapped (**Fig. 4I**). Glutamate and GABA cells were dispersed throughout the PH; there were responders and non-responders throughout the PH without a clear anatomical pattern of connectivity. These results identify functional mPFC-PH connectivity, with mPFC projections synapsing onto both glutamate and GABA neurons but greater connectivity to PH glutamate neurons.

### Stress-Activated PH-Projecting mPFC Circuitry

While mPFC-PH circuit organization has been investigated in males (Myers et al., 2016; reviewed in Schaeuble and Myers, 2022), anatomical connectivity is largely unknown in females. To examine circuit anatomy, male and female Sprague-Dawley rats received microinjections of retrograde-transported AAV construct into the PH and underwent acute restraint stress prior to euthanasia (**Fig. 5A**). Quantification of retrogradely-labeled mPFC neurons revealed a significantly greater density of PH-projecting cells within the female mPFC compared to the male mPFC, with a trend toward a significant interaction between sex and region (2-way ANOVA, main effect of sex F_(1,36)_ = 5.191, *p* = 0.0287, interaction F_(3,36)_ = 2.791, *p* = 0.0543; **Fig. 5B**). The mPFC was subdivided from ventral to dorsal into TT, IL, PL, and cingulate (Cg1) to identify projection neurons within each subregion. Post-hoc analysis revealed a significantly greater number of PH-projecting cells in females compared to males specifically within the TT and IL (Fisher’s LSD test; Male v Female TT, *p* = 0.0244, Male v Female IL, *p* = 0.0258). Stress-induced activation of PH-projecting mPFC neurons was assessed with immunohistochemistry to quantify co-expression of the retrograde-trafficked construct and the immediate early gene, cFos, a marker of recent neural activation (**Fig. 5C, D**). The stress-activation of PH-projecting neurons was dependent on subregion and there was a trend toward a main effect of sex (2-way ANOVA, main effect of region, F_(3,36)_ = 10.72, *p* < 0.0001, main effect of sex, F_(1, 36)_ = 3.878, *p* = 0.0567; **Fig. 5E**). Post-hoc analysis revealed a significantly greater number of stress-activated PH-projecting cells in females compared to males within the PL (Fisher’s LSD test, *p* = 0.0178). When stress activation was normalized to the number of PH-projecting neurons, there was no effect of sex or region (2-way ANOVA, main effect of region F(3, 36) = 1.518, *p* = 0.2263; main effect of sex F(1, 36) = 0.3191, *p* = 0.5757; **Fig. 5F**). This indicates that the *proportion* of PH-projecting mPFC neurons that are activated by stress is similar between the sexes. However, females have a greater overall number of mPFC neurons that project to the PH. Given this difference in mPFC input to the PH, we hypothesized that females would have a greater density of mPFC synapses within the PH.

**Figure 5.**
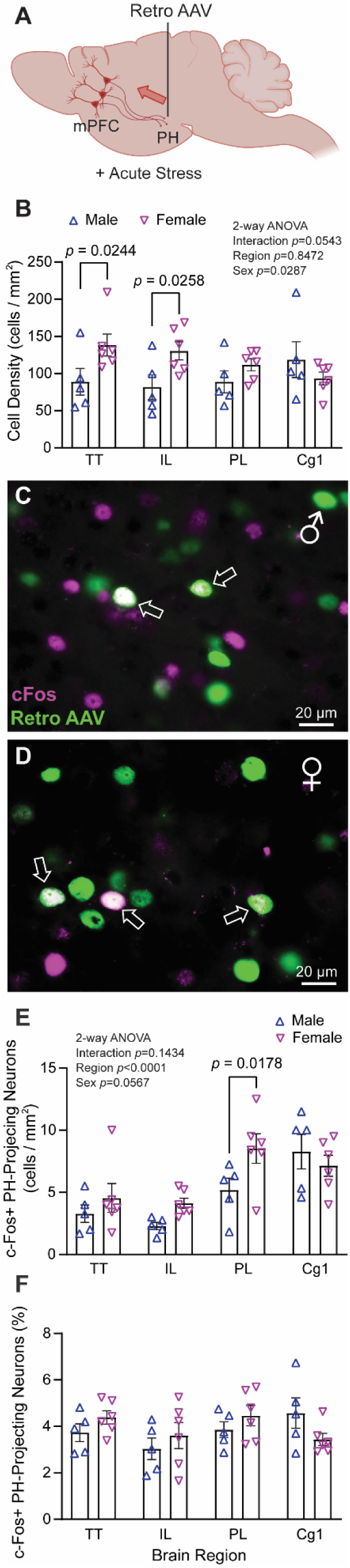
Females have more PH-projecting stress-activated neurons within the mPFC. **A,** Schematic with retro AAV viral injection into PH; **B,** Quantification of retrogradely-labeled neurons within the mPFC; **C,** Representative image of retro AAV and c-Fos within the male mPFC; **D,** Representative image of retro AAV and c-Fos within the female mPFC; **E,** Quantification of retrogradely labeled Fos+ cells within the mPFC; **F,** Percentage of retrogradely labeled cells within the mPFC that are also Fos+. TT, Tenia Tecta; IL, infralimbic region of the prefrontal cortex; PL, prelimbic region of the prefrontal cortex; Cg1, cingulate cortex.

### mPFC Synaptic Terminal Density in the PH

To assess the density of mPFC synapses within the PH, SynaptoTag virus (AAVdj-hSyn-mCherry-GFP-Syb2) was microinjected into the mPFC. SynaptoTag uses the synapsin promoter to induce expression of mCherry in cell bodies and axons of transduced neurons (**Fig. 6A**). Injection area (male: 139.9 ± 36.78 mm^2^, n = 4; female: 190.2 ± 31.63 mm^2^, n = 5) was not statistically different between males and females (unpaired *t* test, t(7) = 1.042, *p* = 0.3320), with similar spread throughout the mPFC (**Fig. 6B-C)**. Conjugation of GFP to synaptobrevin-2 resulted in the expression of GFP at presynaptic terminals of transduced neurons (**Fig 6D**). GFP puncta co-localized with vGluT1 within the PH (**Fig. 6E**), confirming both the terminal expression of GFP and the glutamatergic nature of mPFC-PH projections. Surprisingly, the regional density of GFP within the PH was not different between males and females (unpaired *t* test, t(7) = 0.2201, *p* = 0.821; **Fig. 6G**). Together, these data indicate that despite the larger number of PH-projecting cell-bodies within the female mPFC, the male PH receives similar presynaptic input from the mPFC.

**Figure 6.**
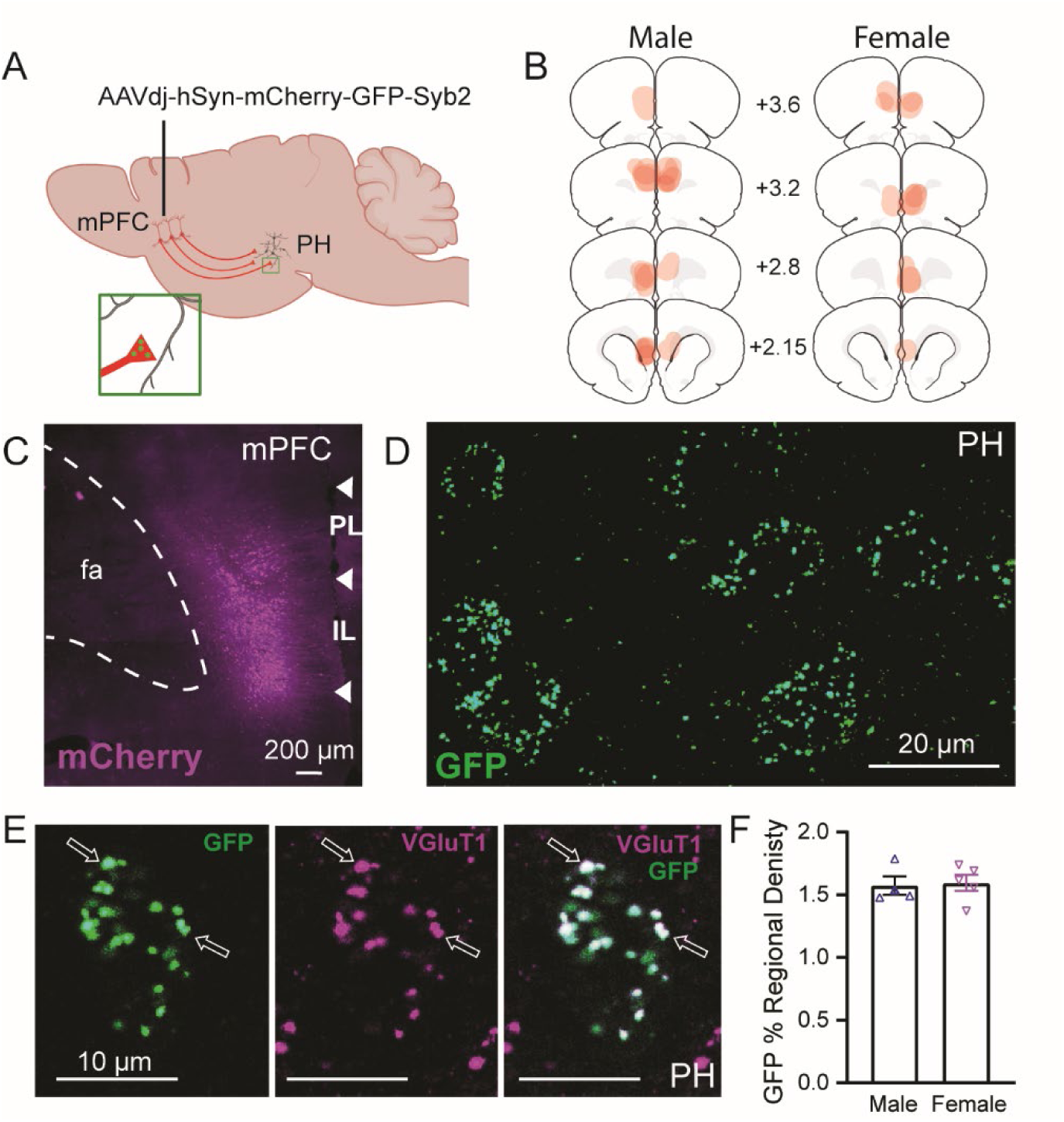
Males and females have similar density of mPFC presynaptic terminals in the PH. AAV-Synaptotag expresses cell-filling mCherry to label soma and axons via the synapsin promoter, GFP conjugated to synaptobrevin-2 labels presynaptic terminals. **A & B**, AAV-Synaptotag was targeted at the mPFC**. C,** Representative image of viral spread within the IL and PL of the mPFC (2.8 mm from bregma; scale bar=200 μm). **D,** Representative SynaptoTag GFP puncta expression in the PH (scale bar = 20um). **E,** Synaptotag GFP puncta colocalization with presynaptic vGluT1 within the PH (scale bar = 10um). **F,** Percent regional density of GFP expression across the rostrocaudal extent of the PH in males and females (N = 4 males, 5 females; *p*=0.9321).

## Discussion

Results presented here detail neurobiological mechanisms for sexually divergent behavioral-homeostatic integration based on mPFC-PH circuit functional organization. We started by investigating neurophysiological properties of PH neurons, as they have not been examined previously. Experiments revealed that sex and estrous phase play a role in membrane and spontaneous firing properties, although neither seem to be the main factors underlying PH neurophysiology. Notably, PH neurons from female rats were more excitable and, although inhibitory tone is observed onto both male and female PH neurons, the frequency of spontaneous inhibitory inputs are greater in males. Functional circuit mapping revealed that in both males and females, glutamatergic mPFC inputs selectively targeted glutamate neurons over GABA neurons within the PH. Both retrograde and anterograde tracing were used to investigate circuit anatomy. Here, a larger ensemble of stress-activated PH-projecting mPFC neurons was present in females; however, the density of mPFC synapses within the PH was not different between sexes. These data outline a neural circuit where females have more stress-processing cortical neurons that target the PH, yet male PH-projecting mPFC neurons likely exhibit greater axonal collateralization or synaptic branching. Collectively, these data identify potential neurobiological mechanisms whereby differences in stimulus processing, pathway organization, inhibitory inputs, and estrous cycle-dependent fluctuations in cellular excitability may culminate in the emergence of differential stress responding between the sexes (**Fig. 7**).

**Figure 7.**
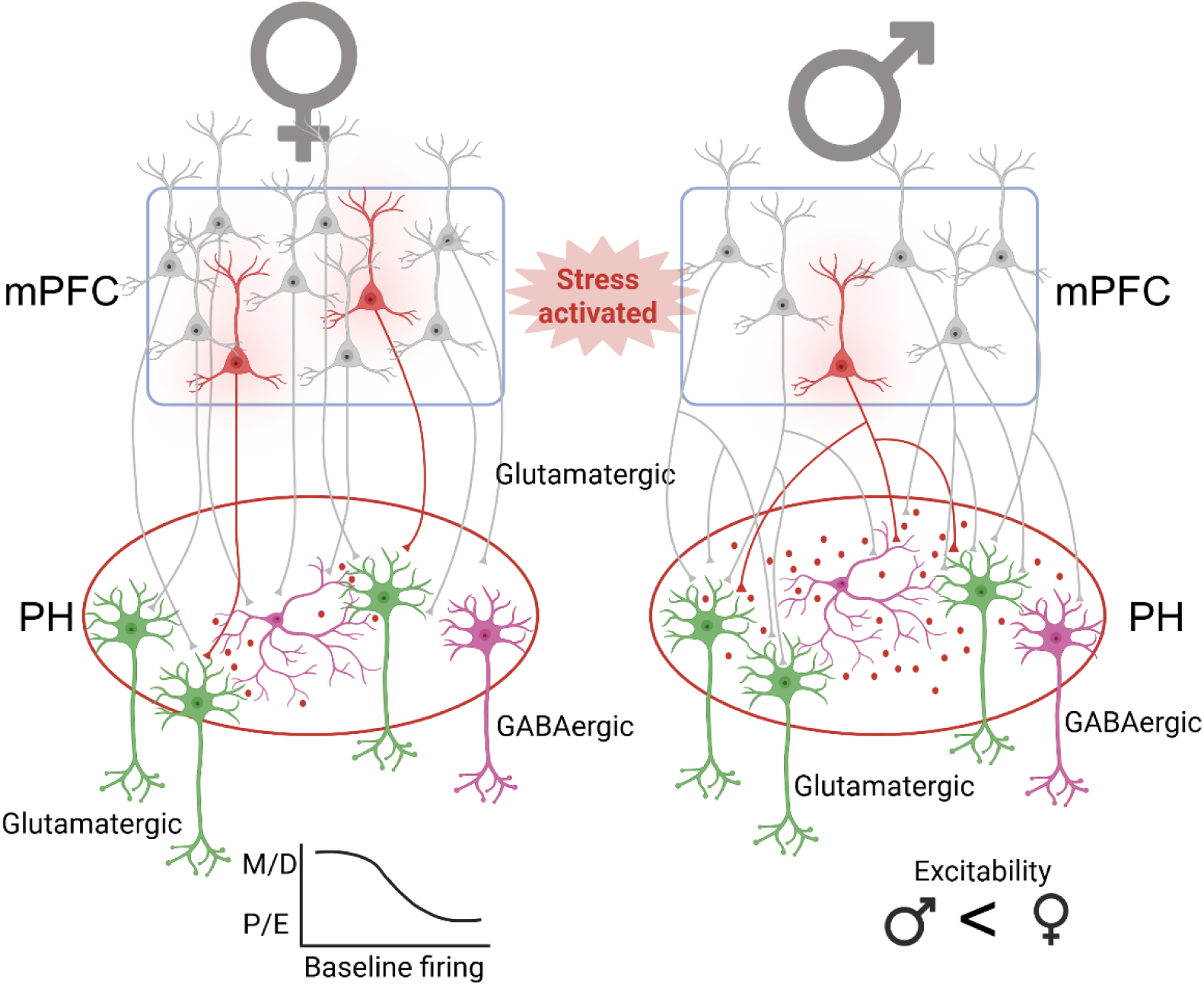
(Summary Figure). Females, compared to males, have significantly more mPFC neurons projecting to the PH with a greater number of stress-activated neurons; however, equal mPFC presynaptic density within the PH in both males and females. Putatively GABAergic and glutamatergic PH neurons receive similar input from the mPFC in males and females. Both male and female PH neurons are under tonic GABAergic inhibition, with the male PH neurons receiving significantly more spontaneous inhibitory input than females. Male and female PH neurons have similar firing properties; although, female neurons have greater excitability and estrous cycle-dependent action potential frequency.

Cell types across the broader hypothalamic synaptic network have been difficult to distinguish physiologically, with the exception of the unique neurophysiological characteristics of neurosecretory hormone-releasing neurons (Renaud and Bourquet, 1991; Tasker and Dudek, 1991; Tasker et al., 2020). Previous histological analyses indicate that the PH is predominantly composed of glutamate and GABA neurons (Abrahamson and Moore, 2001), therefore we combined a Cre-dependent dual reporter virus with a GAD-Cre transgenic rat to visually differentiate, and selectively record from, glutamatergic and GABAergic PH neurons. The GAD-Cre rat has been well validated within the literature (Sharpe et al., 2017; Gibson et al., 2018; Luo et al., 2020; Prasad et al., 2020; Farrell et al., 2022) and > 80% of hypothalamic GAD-expressing cells are Cre+ (Sharpe et al., 2017). Histological analysis was conducted to validate the phenotype of the GFP-expressing PH cells. Examination revealed that nearly 80% of GFP+ cells expressed vGluT2 transcript, while only 12% expressed GAD2 transcript. The GAD+ GFP-expressing neurons likely represent a small population of GABA neurons lacking Cre and/or with failed genetic recombination. We also investigated whether PH dopamine neurons could contribute selectively to either GFP or mCherry populations. We found that TH-positive neurons express GFP and mCherry at approximately similar rates, suggesting portions of this population are GABAergic while some may be glutamatergic. Altogether, the genetic approaches allowed differentiation of PH GFP+ glutamatergic neurons and mCherry+ GABAergic neurons.

Optogenetically-activated mPFC inputs preferentially targeted glutamatergic PH neurons. This experiment combined intersectional colorswitch approaches with CRACM to investigate mPFC connectivity with defined PH postsynaptic targets. Anatomically mapping of synaptic responses identified similar levels of connectivity throughout the PH, congruent with anatomical evidence that the mPFC projections extend throughout the entirety of the male PH (Wood et al., 2019) and identifying a similar connectivity pattern in females. In males, the PH has extensive projections to the PVN which are proposed to contribute to modulation of stress responding (Ulrich-Lai et al., 2011; Myers et al., 2016; Nyhuis et al., 2016). Optogenetic activation of the IL-PH pathway attenuates the physiological stress response in males (Schaeuble et al., 2024) which was hypothesized to occur through increased GABAergic inhibition of the PH outputs to the PVN (Myers et al., 2016). However, here we show that a significantly higher proportion of mPFC inputs synapse onto glutamatergic rather than GABAergic neurons, suggesting that mPFC inputs to the PH could increase PH glutamatergic output. The dense PH projection to peptide-producing cells in the PVN (Myers et al., 2016) might suggest that this mPFC input to the PH could *increase* stress responding; however, the output of these specific glutamatergic neurons is not known. It is also possible that these glutamatergic projections project to the peri-PVN or brainstem and have a more distributed, multi-synaptic impact on physiological stress responses.

Stress-activated mPFC-PH projections have been reported in males (Myers et al., 2016) but had not been investigated in females. Here we show a significantly larger population of PH-projecting mPFC neurons in females compared to males. This effect was driven by higher density of cells in the TT and IL. In males, activation of TT projections to the DMH increase sympathetic activity through DMH projections to sympathetic premotor neurons in the rostral medullary raphe region (Kataoka et al., 2020). However, the role of the TT projection to the PH is not known. Anterograde viruses used throughout these studies were targeted to the IL but some injections spread ventrally into the TT; therefore, some CRACM studies may have also stimulated terminals of TT neurons. The anatomy of mPFC-PH projections has been detailed in males (Myers et al., 2016), with evidence pointing to the PH as an integration center, an intermediary brain region transmitting social and emotional cortical information to the PVN to regulate hypothalamic-pituitary-adrenal axis stress responses (Myers et al., 2016; Nyhuis et al., 2016; Schaeuble and Myers, 2022). In the current study we found more PH-projecting mPFC neurons activated by acute stress in females than males. This was predominantly localized to the PL, which is interesting given evidence that female mice have higher frequency and lower amplitude calcium transients in the PL during immobilization stress (Marin-Blasco et al., 2024). There is evidence that IL and PL have different roles in physiological stress responses (Radley et al., 2006); however, the influence of female stress-activated PL-PH circuitry is not known.

Despite the larger number of PH-projecting cells within the mPFC of females, the density of terminals within the PH of males and females was not different. These data indicate that a smaller number of mPFC neurons can give rise to the same synaptic density in the male, potentially through increased axonal branching or collaterals within this pathway. This sex difference could point to differential mPFC neural processing during stress, whereby females have a larger ensemble of stress-activated, PH-projecting neurons to regulate components of the stress response.

The current results indicate that biological sex and estrous phase play a role in neuronal firing rate and spontaneous action potential properties. Circulating ovarian hormones can modulate neuronal morphology throughout the central nervous system (Woolley et al., 1990; Woolley and McEwen, 1993; Calizo and Flanagan-Cato, 2000; Cooke and Woolley, 2005; Hao et al., 2006) and can modulate spontaneous firing properties (Blume et al., 2017; Willett et al., 2020). Interestingly, the effect of cycle is dependent on the brain region, with different effects of estrous phase within subregions of the basolateral amygdala (Blume et al., 2017). Similar to our findings, others have found that within the nucleus accumbens, sex and estrous cycle play a role in some, but not all, spontaneous firing properties (Proaño et al., 2018; Willett et al., 2020). While circulating estrogens can cross the blood brain barrier and impact hypothalamic firing rates (Yagi, 1973), our receptor expression data suggest that the progesterone receptor is the predominant ovarian steroid receptor in the PH. Thus, the rise in progesterone during P/E may suppress spontaneous firing frequency (Stell et al., 2003; Grassi et al., 2007; Kapur and Joshi, 2021). However, the functional impacts and potential interactions of gonadal hormones within the PH are not known.

These studies advance our understanding of cortical-hypothalamic communication by identifying cell-type and circuit-specific anatomy and neurophysiology. Mechanistically, the combination of anatomical tracing with circuit- and cell-type-specific neurophysiology identifies both structural and functional circuit physiology. The combined anterograde and retrograde tracing delineates anatomical specificities and identifies sexually-divergent circuit organization that could differentially modulate stimulus processing, circuit recruitment, and synaptic signaling during homeostatic challenges. The study of both sexes identifies considerable commonalities and distinctions, highlighting the complexity of cortical-hypothalamic signaling. Signaling components that could substantially modulate functional outcomes display sexual divergence, which may contribute to sex-specific consequences of stress. Together, these findings point to an integrative paradigm where developmental differences in circuit organization, differential cortical processing, as well as sex and hormonal regulation of postsynaptic hypothalamic excitability and inhibitory signaling lead to computational divergence in cortical-hypothalamic communication.

## Acknowledgments

We thank members of the Myers lab for their help throughout these experiments, especially Dr. Derek Schaeuble, Dr. Sebastian Pace, Ema Lukinic, Fallon Stockley, Jake Brown, Helene Duebel, Leo Tyer, Camryn Myers, Tommy Rohr, and Dr. Carley Dearing. We also thank Dr. Bret Smith for advice on preliminary experiments. Schematics were made with Biorender. This project was funded by NHLBI R01-HL173525 (BM) and NHLBI F32-HL172693 (CB).

